# Machine learning surrogate forward models for biomechanical laryngeal control

**DOI:** 10.64898/2026.06.12.731734

**Authors:** Jesús A. Parra, Clara Sorolla, Nicolás F. Quinteros, Emiro J. Ibarra, Gabriel A. Alzamendi, Sean D. Peterson, Hasini R. Weerathunge, Frank H. Guenther, Matías Zañartu

## Abstract

Accurate modeling of laryngeal motor control is key to understanding typical and disordered voice production. However, traditional biomechanical plant models based on ordinary differential equations (ODEs) often involve high computational costs and numerical instabilities, limiting their use in real-time closed-loop control frameworks.

This study evaluates feature-driven machine learning (ML) regressors, specifically Random Forest (RF), Multilayer Perceptron Neural Networks (NN), and Polynomial Regression (PR), as surrogate forward models mapping laryngeal motor inputs to fundamental frequency and sound pressure level. Training data were generated with two biomechanical vocal fold models: the extended body-cover and the triangular body-cover.

Results demonstrate that ML surrogates reduce execution times from seconds to milliseconds (e.g., 2 ms for PR), enabling stable real-time tracking via inverse Jacobian control. While RF provides the highest accuracy, NN and PR offer smoother control signals and smaller memory footprints. A practical performance threshold was identified near N = 1,000 training samples, below which accuracy degraded substantially when models were trained from scratch.

These findings support ML surrogates as efficient and adaptable alternatives to direct numerical simulation, providing a foundation for future subject-specific modeling through transfer learning in data-limited clinical scenarios.

## I. INTRODUCTION

Biomechanical control of the larynx plays a central role in voice production by regulating the motion of the vocal folds (VFs), enabling precise control of key acoustic parameters such as the fundamental frequency (f*_o_*) and sound pressure level (SPL). These outputs arise from the coordinated interaction between subglottal pressure (P*_S_*) and intrinsic laryngeal muscles (ILMs) activation, primarily the cricothyroid (a*_CT_*) and thyroarytenoid (a*_T A_*) muscles, which together govern pitch, loudness, and phonatory stability, all of which are essential for intelligible speech (Chhetri and Neubauer, 2015; Zhang, 2016). Beyond normal voice production, alterations in this biomechanical control are closely associated with voice disorders, highlighting the need for physiologically grounded models that can capture the relationship between motor control inputs and resulting acoustic outputs (Gomez *et al*., 2007; Hillman *et al*., 2020). Although the ultimate objective in phonatory control is to solve the inverse problem, this process fundamentally depends on accurate forward mappings. To this end, the present work evaluates feature-driven machine learning (ML) regressors as foundational surrogate forward models for laryngeal control, providing a necessary step toward enabling control-oriented modeling of phonation.

In the context of speech motor control, laryngeal motor commands are typically embedded within a framework aimed at generating desired phonatory behaviors. Speech control models have typically and primarily focused on the articulatory control of vocal tract structures (Kröger *et al*., 2009; Perrier *et al*., 2006; Saltzman and Munhall, 1989; Tourville and Guenther, 2011). More recently, extensions such as LaDIVA (Weerathunge *et al*., 2022) have incorporated laryngeal biomechanical control, where ILMs activation and P*_S_* are mapped to acoustic outputs (f*_o_* and SPL) through biomechanical models of VFs dynamics. In parallel, recent work has explored the use of deep neural networks within laryngeal neuromuscular control architectures, replacing traditional controllers with learned feedforward and feedback mappings that integrate both acoustic and somatosensory targets while retaining a physics-based VFs model as the controlled plant (Palaparthi *et al*., 2024). Within this context, phonation can be formulated as a tracking problem, where desired acoustic targets are achieved by iteratively adjusting motor control inputs. A forward model plays a central role in this process by predicting the acoustic output resulting from candidate control inputs, enabling feedback controllers to minimize the error between desired and produced signals. Consequently, the performance of the forward mapping directly impacts the accuracy and stability of control-oriented simulations.

Modeling laryngeal control remains challenging due to the nonlinear and dynamic behavior of the VFs. Lumped-element models, such as the extended body-cover model (ext-BCM) (Zañartu *et al*., 2014) and the triangular body-cover model (TBCM) (Alzamendi *et al*., 2022), provide a reasonable balance between interpretability and computational efficiency. However, full-phonatory biomechanical models require the numerical solution of large and complex systems of differential equations (ODEs). In practice, simulating a single steady-state condition can take approximately few seconds of real computational time. Within a feedback control loop operating in tight iterative windows (e.g., 5 ms), this cost can scale to minutes or hours for just one second of speech, rendering direct ODE integration restrictive. Furthermore, these models exhibit mathematically unstable regions, such as extreme a*_CT_*-a*_TA_* imbalances, where standard periodic metrics like f*_o_* fail to converge, creating “holes” or undefined values that can cause gradient-based controllers to fail (Weerathunge *et al*., 2022).

Data-driven approaches based on ML have shown strong potential for approximating nonlinear relationships in voice production, particularly in inverse problems such as estimating P*_S_* or muscle activations from acoustic features (Donhauser *et al*., 2024; Ibarra *et al*., 2021; Marks *et al*., 2020). In these settings, latent physiological variables are inferred from observable non-invasive signals, such as microphone recordings, which act as reliable proxies for laryngeal dynamics. However, existing ML approaches have predominantly focused on parameter estimation rather than modeling the forward mapping within control-oriented frameworks. Likewise, although recent neural control architectures have incorporated ML-based controllers, they still rely on computationally expensive biomechanical plants for forward simulation (Palaparthi *et al*., 2024). Related efforts in speech motor control have also introduced modernized implementations of the original DIVA framework, primarily aimed at improving extensibility and integration with contemporary ML tools while retaining physics-based synthesis components (Kinahan *et al*., 2023). In particular, the suitability of ML-based surrogate models in terms of accuracy, computational efficiency, and stability under iterative conditions has not been systematically evaluated. This gap limits the integration of ML models into biomechanical control architectures.

To address these limitations, ML-based surrogate models offer a fast and continuous alternative for approximating the mapping between laryngeal control inputs and acoustic outputs. By learning from data generated by biomechanical models, these approaches can significantly reduce computational costs (e.g., reducing evaluation times from seconds to milliseconds) while preserving the essential physics. In contrast to numerical solvers, ML surrogates provide smoother approximations across regions where conventional models exhibit instability. Lookup Table (LUT)-based approaches may also hinder subject-specific personalization and adaptation to unseen phonatory regimes. Furthermore, while personalizing traditional ODE models requires solving ill-posed optimization problems to adjust hidden anatomical hyperparameters, an ML surrogate provides a flexible architecture suited for data-driven fine-tuning via transfer learning (TL).

This study aims to develop and systematically evaluate ML-based surrogate forward mappings for laryngeal biomechanical control, comparing Random Forest (RF), Multilayer Perceptron (NN), and Polynomial Regression (PR) architectures. Feature-driven surrogates are especially attractive in low-data scenarios due to their lower training burden and greater interpretability. The proposed framework is assessed with emphasis on accuracy, computational efficiency, and its suitability for tracking phonatory targets within a closed-loop control framework. Additionally, by analyzing model performance under varying data conditions between two lumped-element models, this work establishes a foundational baseline for future subject-specific adaptation and clinical translation.

### II. MATERIALS AND METHODS

This section details the development and validation of the feature-driven ML surrogate forward models. As discussed in Sec. I, the primary goal is to establish a foundational framework that overcomes the computational bottlenecks and instabilities of numerical ODE solvers, enabling real-time laryngeal motor control simulations. To address this objective, the methodology is organized as follows: (i) we first define the conceptual laryngeal control framework, identifying the roles of the planner, controller, and plant within a closed-loop tracking system (section II A); (ii) we then describe the biomechanical VFs models and the structural architecture of the numerical plant used to generate the synthetic datasets (section II B); (iii) we detail the specific ML architectures and training procedures selected to approximate these large and complex mappings (section II C); and (iv) we present the feedback control scheme and the pitch and loudness glide exercises designed to evaluate the dynamic performance of the surrogates (section II D).

### A. Laryngeal control framework

Speech motor control models are typically conceptualized as closed-loop systems comprising three fundamental components: a planner, a controller, and a plant (Guenther, 2016; Parrell *et al*., 2019). As illustrated in Figure 1(a), the **planner** establishes the desired sensory targets (y*_T_*), which are compared against the current system output (y(t)) to generate an acoustic error signal (ɛ(t)). The **controller** then processes this error to compute the necessary motor commands (x(t)) required to drive the plant toward the target. In the context of laryngeal biomechanics, the **plant** represents the forward transformation from physiological motor inputs to acoustic outputs.

**FIG. 1.**
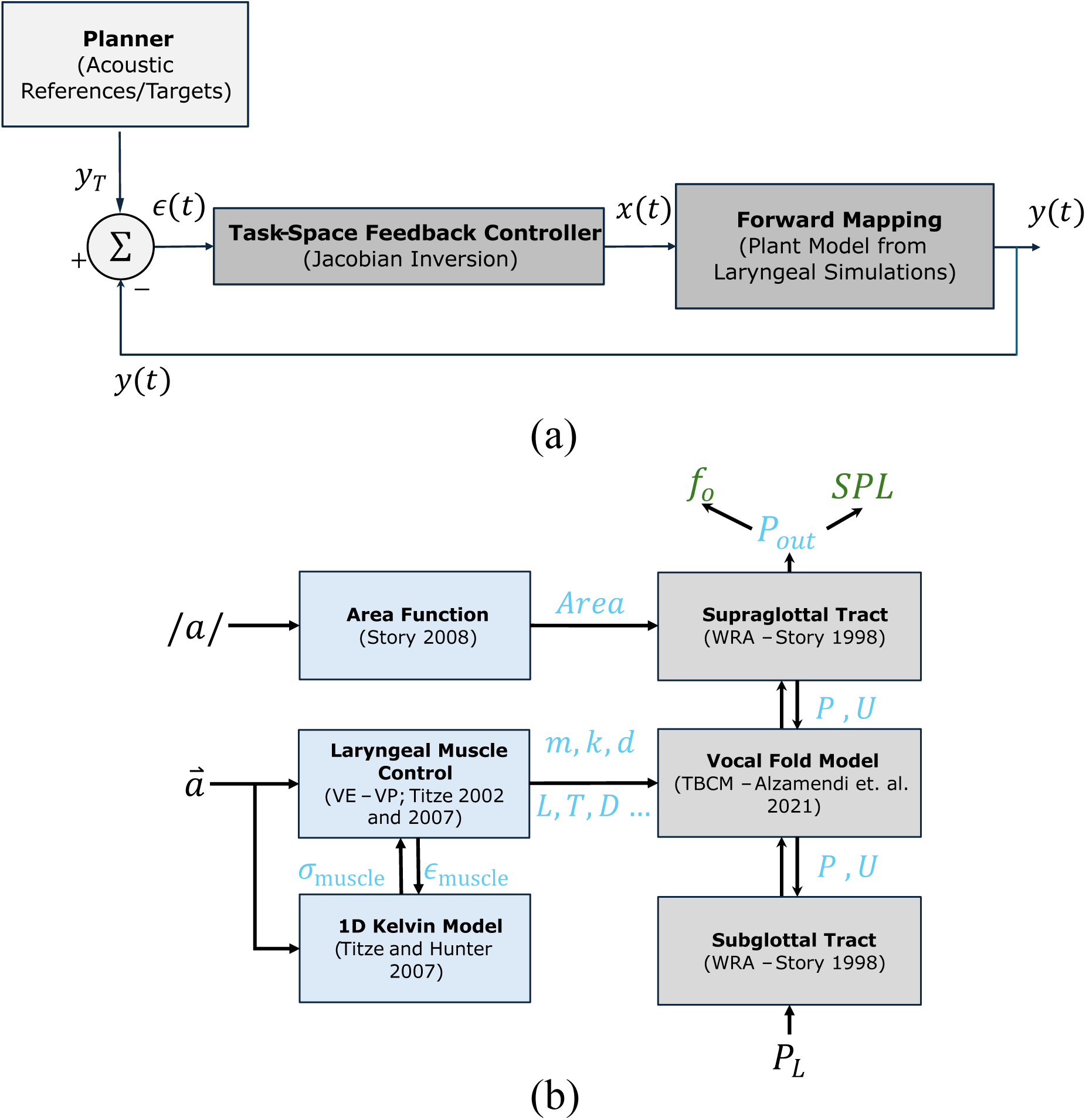
Overview of the control and simulation framework: (a) Closed-loop laryngeal control scheme illustrating the tracking process via an inverse Jacobian controller. (b) Numerical plant used for dataset generation, showing the transformation of motor inputs 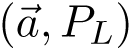 into tissue mechanical properties (1D Kelvin model), the interaction with the core ODE vocal fold model, and acoustic wave propagation (WRA) to yield the radiated sound pressure (P*_out_*), f*_o_*, and SPL.

In this study, the motor commands x(t) are defined by the activation of the ILMs (a*_CT_*, a*_T A_*) and P*_S_*, while the acoustic outputs y(t) correspond to f*_o_* and SPL. Effective tracking of these acoustic targets requires the controller to iteratively solve an inverse problem, which in turn depends on a robust and continuous forward mapping within the plant. While this mapping can be achieved through direct numerical integration of laryngeal differential equations (detailed in section II B), the computational bottlenecks and mathematical instabilities discussed in section I make such an approach restrictive for the repeated evaluations required in a closed-loop setting.

Consequently, the **Forward Mapping** block in Figure 1(a) is represented in this work by different ML architectures acting as surrogate models. These regressors provide a fast, data-driven approximation of the vocal function, enabling the real-time computation of acoustic outputs from motor inputs. To assess the viability of these ML-based plant representations, we design dynamic control exercises focused on tracking phonatory gestures (pitch and loudness glides), analyzing how the choice of regressor influences the stability and accuracy of the feedback correction process.

### B. Vocal fold models and dataset generation

Lumped-element biomechanical models provide computationally efficient representations of the dynamics of VFs by using mass-spring-damper systems. These models capture key biomechanical interactions while maintaining a balance between physiological accuracy and computational feasibility. In this study, we selected two lumped-element VFs models, ext-BCM and TBCM, that capture different aspects of phonation dynamics: the ext-BCM provides a simplified yet physiologically grounded representation of VFs motion, while the TBCM extends this framework by incorporating a more anatomically designed configuration and a physiologically representative description of activation of the ILMs. These models were selected to evaluate regressor behavior across a controlled range of biomechanical complexity while preserving a common body–cover, three-mass architectural framework. The ext-BCM (Zañartu *et al*., 2014) implements empirical body–cover rules (Titze and Story, 2002) to map ILMs activation to effective tissue parameters. In contrast, the TBCM (Alzamendi *et al*., 2022) incorporates a triangular VFs geometry and explicit stress–strain relations that more directly reflect muscle-driven tissue mechanics (Titze and Hunter, 2007).

The structural architecture of the numerical plant is illustrated in Figure 1 (b). The system is driven by motor command inputs, including the vector of intrinsic muscle activations (a⃗) and lung pressure (P*_L_*). Within the laryngeal muscle control block, a 1D Kelvin model (Titze and Hunter, 2007) determines the active and passive muscle stress (σ*_muscle_*) and strain (ɛ*_muscle_*) to specify the effective mass (m), stiffness (k), and damping (d) parameters, as well as the geometric dimensions (length L, thickness T, and depth D) of the VFs. These mechanical properties serve as parameters for the core ODE VFs model, which interacts via pressure (P) and airflow (U) with the subglottal and supraglottal tracts. The supraglottal tract is configured with a fixed area function for the vowel /a/ (Story, 2008), and both tracts utilize a Wave Reflection Analog (WRA) (Story and Titze, 1995) to account for source-filter interaction and acoustic propagation. The final output is the radiated acoustic pressure (P*_out_*), from which the target features f*_o_* and SPL are extracted.

Simulations were performed for sustained (quasi-steady state) vowel phonation conditions, systematically varying the motor inputs (P*_S_*, a*_CT_*, a*_T A_*) in physiologically relevant ranges. For each data point, these input parameters were held constant throughout the simulated interval. The models were initialized and allowed to run for 0.5 s prior to any feature extraction. Given the simulated phonatory frequency range (approximately 100-300 Hz), this 0.5 s initialization corresponds to around 50 to 150 glottal cycles, which sufficient to allow all initial transients to decay and the dynamical system to reach a stable limit-cycle oscillation. Acoustic features were then computed from the final 50 ms of the signal. Specifically, f*_o_* was derived from the simulated glottal airflow waveform (U in Figure 1 b), as it represents the source excitation signal and is robust to vocal tract filtering effects. SPL was computed from the radiated sound pressure signal in front of the lips (P*_out_* in Figure 1 1b) as the unweighted overall SPL: SPL = 20 log_10_ (p*_rms_*/p*_ref_*), where p*_rms_* is the root-mean-square acoustic pressure, and p*_ref_* = 20 µPa.

The ext-BCM dataset was generated using the VFs model by Zañartu *et al*. (2014), which is implemented as a plant to be controlled in the LaDIVA model through an LUT (Weerathunge *et al*., 2022). This VFs model extends the classical body-cover model originally proposed by Story and Titze (1995), differentiating between the body and cover layers of the VFs while incorporating a fixed posterior glottal opening. In addition, it models neuromuscular activation of key ILMs: CT, TA, and LCA, along with P*_S_*. These activations influence VFs stiffness and geometry (Titze and Story, 2002), with muscle activation levels normalized between 0 (rest) and 1 (maximum contraction). For dataset generation, LCA activation was fixed at 0.5, while a*_CT_* and a*_T A_* were varied from 0 to 1 in 0.01 increments, and P*_S_* was adjusted from 10 to 2010 Pa in 25 Pa increments. This resulted in a comprehensive dataset consisting of about 800,000 data points.

The TBCM dataset was generated using the TBCM model described in Alzamendi *et al*. (2022), which extends the body-cover framework by incorporating a triangular VFs configuration (Galindo *et al*., 2017). This model accounts for the vocal process variations in VFs motion, providing a more detailed characterization of phonatory biomechanics. It includes the five activations of the ILMs which interact to regulate VFs stiffness, mass distribution, and vocal posture (Titze and Hunter, 2007; Titze and Story, 2002), and a more detailed ILMs stress-strain relation with a muscle Kelvin model (Titze and Hunter, 2007). Simulated dataset for typical phonation was obtained by systematically varying a*_CT_* and a*_T A_* from 0 to 1 in 0.025 increments and P*_S_* from 500 to 2,000 Pa in 50 Pa increments. To ensure complete glottal closure, LCA and IA activations were fixed at 0.5, while PCA activation was set to 0. All simulations were conducted using a male phonatory system configuration, yielding approximately 50,000 data points.

The dataset sizes were intentionally disparate (approx. 800,000 samples for ext-BCM vs. 50,000 for TBCM). This setup serves as a preliminary exercise to observe model behavior across different data regimes, emulating the restricted data availability typically found in subject-specific or high-fidelity clinical scenarios. Table I details the input scheme adopted for each model, summarizing the specific muscle activation ranges and the resulting dataset dimensions; for this study, anatomical parameters were fixed to represent a standard male laryngeal geometry to establish a methodological baseline.

**TABLE I.**
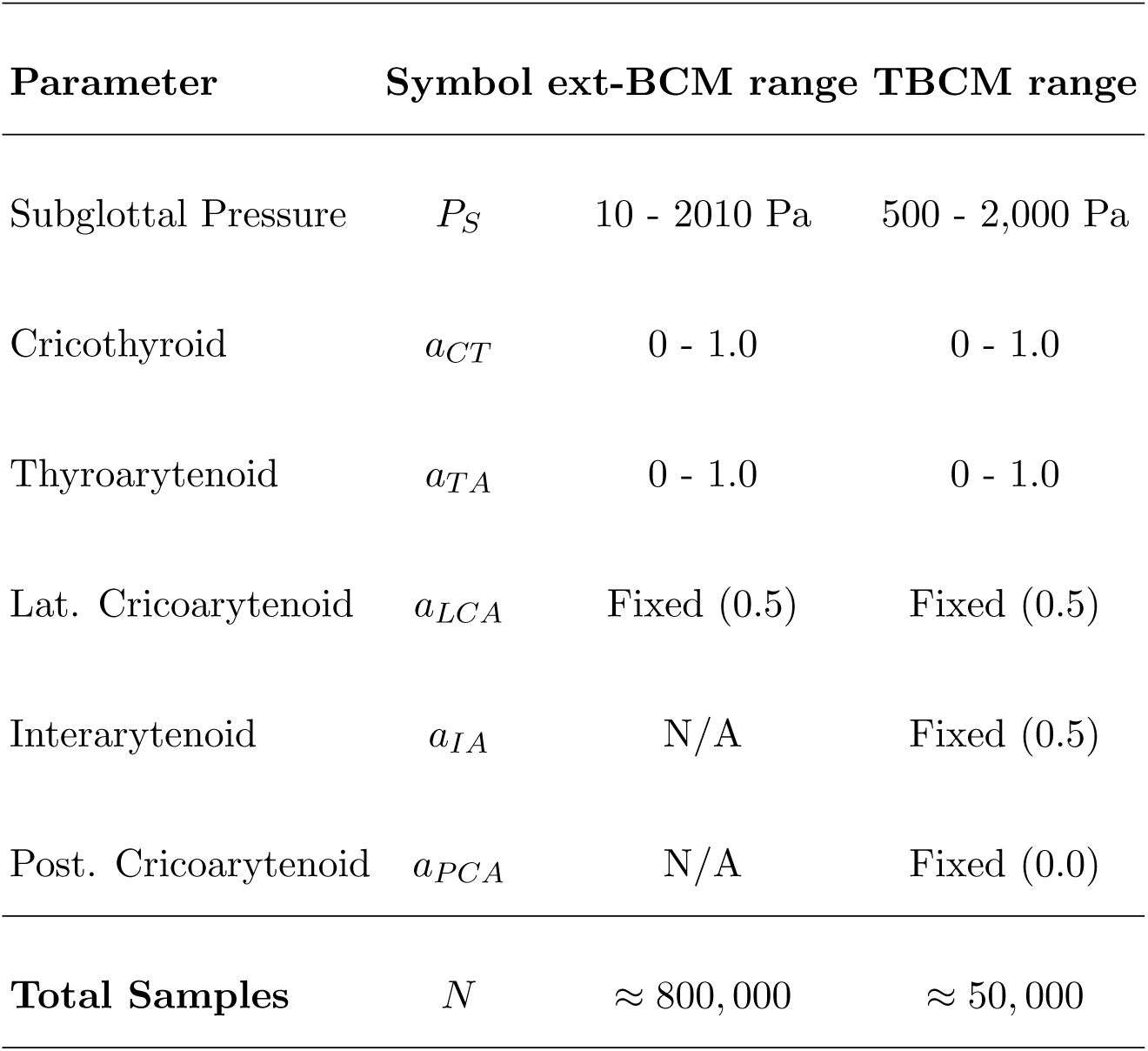
Summary of input parameters, ranges, and fixed values for the generated datasets. Output parameters for both models were Fundamental Frequency (f*_o_*) and Sound Pressure Level (SPL).

| Parameter | Symbol | ext-BCM range | TBCM range |
| --- | --- | --- | --- |
| Subglottal Pressure | $P_S$ | 10 - 2010 Pa | 500 - 2,000 Pa |
| Cricothyroid | $a_{CT}$ | 0 - 1.0 | 0 - 1.0 |
| Thyroarytenoid | $a_{TA}$ | 0 - 1.0 | 0 - 1.0 |
| Lat. Cricoarytenoid | $a_{LCA}$ | Fixed (0.5) | Fixed (0.5) |
| Interarytenoid | $a_{IA}$ | N/A | Fixed (0.5) |
| Post. Cricoarytenoid | $a_{PCA}$ | N/A | Fixed (0.0) |
| **Total Samples** | $N$ | $\approx 800,000$ | $\approx 50,000$ |

### C. Machine learning regressors

To implement the forward mapping, we assessed the performance of five ML regressors selected to represent three complementary modeling families. This selection strategy was designed to explore the tradeoffs between predictive accuracy, output smoothness, interpretability, and computational cost. Implementation, training, and evaluation tasks for all the considered models were performed in Python (version 3.9) using skikit-learn (version 1.6) and Keras libraries. All models were trained on a workstation equipped with an Intel Core i5-10400F CPU @ 2.9 GHz, 128 GB of RAM, and an NVIDIA RTX 3050 GPU.

Polynomial Regression (PR) was selected as an interpretable baseline (Rudin, 2019). It extends linear regression by fitting polynomial functions to input variables, allowing it to capture low-order non-linearities while maintaining a lightweight computational footprint suitable for rapid evaluation. Tree-Based Methods, including Random Forest (RF), Gradient Boosting (GB), and Extreme Gradient Boosting (XGB), were employed to represent ensemble learning techniques (Grinsztajn *et al*., 2022). These models construct multiple decision tree structures to capture complex, tabular non-linearities and variable interactions, offering high accuracy and robustness against outliers. Finally, a Neural Network (NN) regressor was implemented as a Multilayer Perceptron (MLP). As a universal function approximator, the NN is capable of modeling high-dimensional non-linear relationships (Hunt *et al*., 1992). Crucially for this application, it provides smooth continuous mappings, a desirable property for gradient-based control frameworks.

The tree-based regressors were implemented with the “MultiOutputRegressor” strategy, which allows fitting a separate model for each target variable in parallel, and training using their default configurations. The NN regressor consists of an input layer with three neurons, three hidden layers with 128 neurons each, and an output layer with two neurons, inspired by previous regression work (Ibarra *et al*., 2021). Each hidden layer employs a Rectified Linear Unit (ReLU) activation function and incorporates a dropout layer of 0.1 to mitigate overfitting. The PR was implemented using “PolynomialFeatures” function, which generates a new feature matrix consisting of all polynomial combinations of the input features up to a specified degree, after which a linear regression model is trained on the expanded feature set.

For all regressors, the inputs were a*_CT_*, a*_T A_*, and P*_S_*, while the outputs were f*_o_* and SPL. The dataset derived from the ext-BCM model was used for training, after min-max normalization. The training procedure was organized as follows. First, the data was randomly split into 80% training and 20% testing to perform an initial evaluation of the three tree-based regressors (RF, GB, XGB). This allowed the selection of the most promising (accurate) tree-based method to be studied in detail. Second, hyperparameter tuning was conducted for the selected tree-based model, the PR, and the NN, following a 5-fold cross-validation technique on the entire dataset, ensuring that every data subset was used for both training and testing in a rotational manner. Importantly, the validation folds acted as independent test subsets that did not participate in model fitting, providing unbiased estimates of generalization performance.

For the PR, the polynomial degree was varied from 1 to 15. For the tree-based model, the number of estimators was varied between 50 and 200, and the maximum depth between 20 and 60. Hyperparameter tuning was performed using the “GridSearchCV” method from Scikit-learn, employing 5-fold cross-validation to identify the optimal configuration for each regressor (Pedregosa *et al*., 2011). For the NN regressor, the same 5-fold strategy was applied: the validation fold in each split was used exclusively for performance evaluation, without being incorporated into gradient descent optimization or early stopping criteria.

Model performance was assessed using mean absolute error (MAE), root mean square error (RMSE), and the coefficient of determination (R^2^). Errors were computed between the regressor-predicted values of f*_o_* and SPL and the corresponding ground-truth values obtained from the ext-BCM and TBCM simulations. For each regressor, the best performing configuration over all folds was selected, reflecting the model that most accurately captured the input–to–output mapping.

Approximate training durations were 19 s and 4 s for PR, 370 s and 70 s for NN, and 106 s and 16 s for RF. These timing differences reflect the computational complexity associated with training each architecture. However, because training is a one-time offline process, the true computational advantage of the ML framework relies on its inference (execution) time during the closed-loop control scheme, which is orders of magnitude faster than direct ODE integration (as detailed in Section III).

Finally, the best-performing ML regressors trained on the ext-BCM dataset were retrained from scratch using the TBCM dataset to evaluate its adaptability across different laryngeal representations. Keeping the same 5-fold cross-validation strategy, the regressor performances were evaluated using MAE, RMSE, and R^2^. This step was included to test the capacity of the regressors to generalize across datasets of different magnitudes, while the use of synthetic data in the first stage provides a robust foundation for subsequent fine-tuning with more limited experimental datasets. This experiment can be seen as a first synthetic example of TL, designed to assess how the three regressors handle both reduced data availability and a change in variable relationships. While a slight drop in performance is expected, the results remain relevant as an indicator of their robustness.

### D. Feedback error using inverse Jacobian method for reference tracking

Following the laryngeal control framework described in Sec. II A, this section details the implementation of the **Feedback Controller** and the dynamic tracking exercises used to evaluate the ML-based plant. The controller’s role is to translate the estimated sensory error ɛ(t) = y*_T_* (t) − y(t) into iterative updates for the motor command vector x(t) = [a*_CT_*, a*_T A_*, P*_S_*]*^T^*. Following the LaDIVA framework (Weerathunge *et al*., 2022), the feedback controller translates estimated auditory error into the motor command space using a linear approximation of the forward mapping. The update rule for adjusting the input state x is given by:

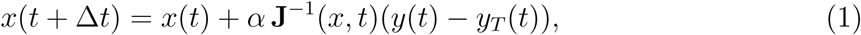

where α = 0.1 is the step size and **J***^−^*^1^ is the inverse Jacobian of the forward mapping, computed using a modified Moore-Penrose pseudoinverse with damped least squares penalization:

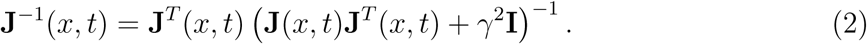

with the damping parameter set to γ = 0.1. The numerical derivatives for the Jacobian were evaluated using a step size of 0.02 for the normalized inputs.

To make the closed-loop mechanics (illustrated in Figure 1 (a)) explicit, the tracking process operates iteratively as follows: (1) The *Planner* defines the desired acoustic targets at a given time step; (2) an error is calculated between this target and the current acoustic output; (3) the *Feedback Controller* (Inverse Jacobian) computes the necessary physiological corrections (Δa*_CT_*, Δa*_T A_*, ΔP*_S_*) to reduce the acoustic error; (4) these updated physiological values are fed directly into the *Forward Mapping* (the ML regressor); and (5) the ML regressor computes the new f*_o_* and SPL; and the cycle repeats for the next time step.

As established in the methodological roadmap, we integrated this controller with the ML surrogates to perform two types of sensory reference targets (y*_T_*): a pitch glide and a loudness glide. The pitch glide consists of a quadratic increase in f*_o_* (from 150 Hz to 300 Hz) at a constant 70 dB SPL, while the loudness glide involves a linear SPL increase (65 to 85 dB for ext-BCM; 75 to 110 dB for TBCM) at a fixed f*_o_* of 200 Hz.

These reference gestures serve as a controlled benchmark to evaluate the capability of the ML-based plants to generate smooth and continuous output trajectories y(t) in response to the controller’s commands. By mimicking time-varying laryngeal gestures, this exercise assesses whether the accuracy and stability of the surrogate mappings are maintained under the iterative, closed-loop conditions required for speech motor control simulations.

## III. RESULTS

This section presents the evaluation of ML regressors for forward mapping and reference tracking using both the ext-BCM and TBCM datasets. The simulations are structured as follows. First, the performance of tree-based regressors is evaluated, and optimized regressors from different model families are assessed. Then, the predictions of each optimized ML regressor are compared to the original dataset to evaluate accuracy, estimation time (as a proxy for computational complexity), and model size (memory footprint). Finally, the performance of each optimized regressor to track reference acoustic targets for pitch and loudness glide gestures within a control scheme is addressed.

### A. Evaluation of machine learning regressors for forward mapping and reference tracking using the ext-BCM dataset

The initial evaluation was performed using the ext-BCM dataset, which served as the primary training set for the regressors. The results for the tree-based regressors are presented in Table II, comparing the MAE, RMSE, and R^2^ scores of the estimated f*_o_* and SPL. The RF and XGB regressors achieved the highest R^2^ scores of 0.99. Among the other metrics, RF stood out, showing the lowest estimation errors, with MAE and RMSE below 1 Hz for f*_o_* and below 0.1 dB for SPL. Therefore, RF was selected as the best tree-based regressor for further analysis.

**TABLE II.** Error metrics for initial direct mapping fitting of ext-BCM f*_o_* and SPL for tree-based methods. RF: Random Forest, GB:Gradient Boosting, XGB: Extreme Gradient Boosting.

| Model Regressor | $f_o$ | | | $SPL$ | | |
| --- | --- | --- | --- | --- | --- | --- |
| | $R^2$ | MAE (Hz) | RMSE (Hz) | $R^2$ | MAE (dB) | RMSE (dB) |
| **RF** | **0.999** | **0.355** | **0.578** | **0.999** | **0.053** | **0.074** |
| GB | 0.985 | 8.907 | 12.656 | 0.984 | 0.845 | 1.033 |
| XGB | 0.999 | 1.368 | 1.854 | 0.999 | 0.084 | 0.120 |

Table III summarizes the error metrics of the optimized regressors, including RF, NN, and PR models. The regressors achieved excellent R^2^ scores, exceeding 0.998 in all cases. The optimized RF regressor exhibited the lowest estimation errors, with less than 0.5 Hz for f*_o_* and less than 0.05 dB for SPL. Note that the error metrics for SPL were computed on a logarithmic scale, expressed in dB re 20 µPa. Therefore, the reported MAE and RMSE quantify the average absolute and quadratic errors directly in decibels, maintaining physical interpretability since both predictions and ground-truth targets share the exact same reference pressure.

**TABLE III.** Error metrics after cross-validation and hyper-parameter fitting for direct mapping fitting of ext-BCM f*_o_* and SPL for Random Forest (RF), Neural Network (NN) and Polynomial Regressor (PR).

| | $f_o$ | | | $SPL$ | | |
| --- | --- | --- | --- | --- | --- | --- |
| | $R^2$ | MAE (Hz) | RMSE (Hz) | $R^2$ | MAE (dB) | RMSE (dB) |
| RF | 0.999 | 0.172 | 0.473 | 0.999 | 0.023 | 0.039 |
| NN | 0.999 | 0.878 | 1.491 | 0.998 | 0.357 | 0.513 |
| PR | 0.999 | 0.531 | 1.424 | 0.999 | 0.183 | 0.321 |

In terms of f*_o_*, the NN and RF regressors had similar RMSE values for f*_o_*, approximately 1.5 Hz, while for the remaining metrics, PR showed a slight improvement over NN.

To further compare the regressors, Figure 2 illustrates the inference (execution) time as a function of sample size. It shows the mean inference time over 10 runs for different sample sizes, displayed on a logarithmic scale for better visualization. As discussed in Section I, direct numerical integration of the ODE system takes approximately 6 s per sample. In stark contrast, all ML regressors operate in the millisecond or sub-millisecond range. PR was the fastest method, yielding a half and one order of magnitude reduction, respectively, in computational time compared to RF and NN regressors. Specifically, PR required 2×10*^−^*^3^ s (∼2 ms) for a single sample (a 3-component vector) and 4 s for a dataset of size 10^6^. On the other hand, NN initially required the longest time to generate an estimate (5 × 10*^−^*^2^ s); however, for sample sizes above one thousand samples, it overtook RF in efficiency. At 10^6^ samples, NN required 44 s, compared to RF’s 67 s. Sample size can be an important factor when performing multiple parallel computations.

**FIG. 2.**
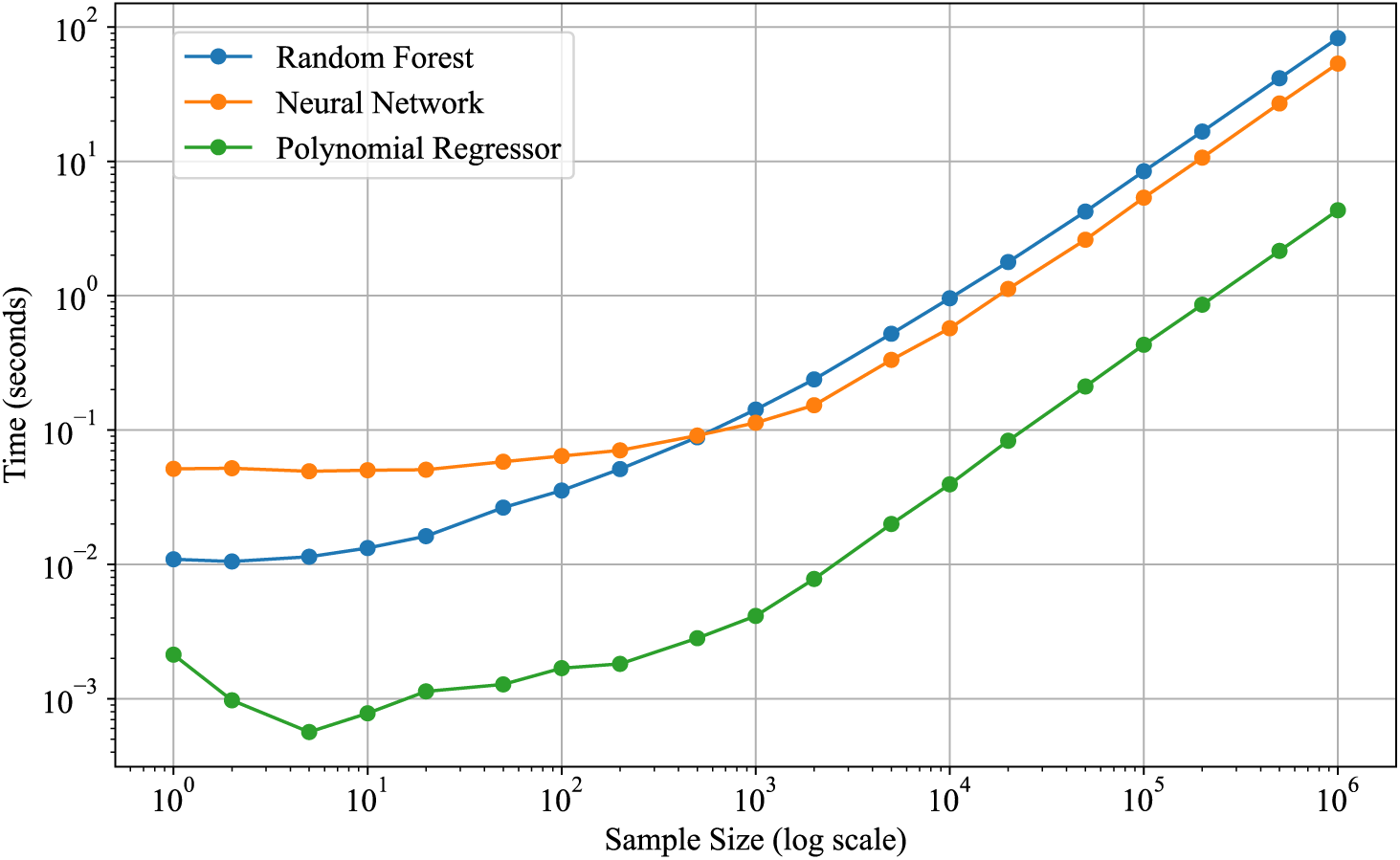
Inference (execution) time vs sample size for the different ext-BCM-based regressors, using Random Forest (RF), Neural Network (NN) and Polynomial Regressor (PR).

The considered regressors were also analyzed in terms of model size (in KB), reflecting their complexity and storage requirements. The lightest models were PR (degree 12, 16 KB) and NN (3 hidden layers, 128 neurons, 438 KB). In contrast, RF (100 estimators, maximum depth of 40) required the largest storage, occupying 1.3 GB, as a result of the large number of conditional (’if’) operations required for the estimation process.

To provide a graphical representation of the regressor estimations, heatmaps (Figure 3) were used to analyze the relationship between muscle activation and the predicted values of f*_o_* and SPL. For this analysis, P*_S_* was fixed at 1 kPa, a value approximately associated with loud conversational speech (Titze, 2000). The physiological rationale for computing these maps at 1 kPa was to assess whether the ML regressors reproduce the expected biomechanics under a representative modal phonation regime at moderate-to-high vocal intensity. Fixing the driving pressure allows us to isolate and visualize the highly non-linear interactions of laryngeal tension (a*_CT_* and a*_T A_*) on f*_o_* and SPL. The results show that frequency increases with CT muscle activation, with a more curved relationship at low frequencies and a linear relationship at higher frequencies. Notably, all regressors produced highly similar estimations, which can be attributed to the flexibility for learning the latent, non-linear dependencies from a large number of training samples. This aspect will be contrasted in the following section when evaluating regressors with the TBCM dataset.

**FIG. 3.**
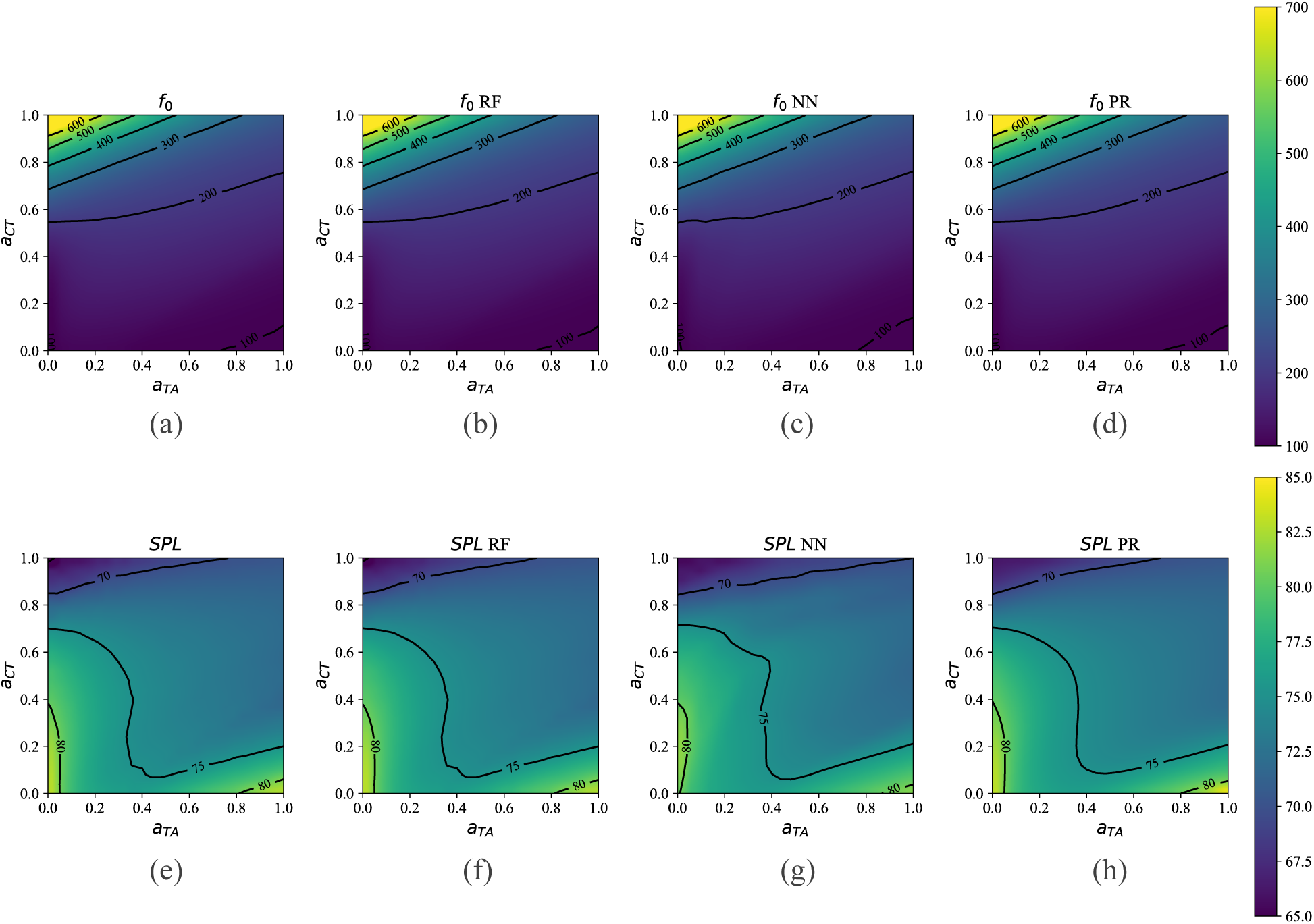
MAP of f*_o_* (a-d) in Hz and SPL (e-h) in dB for the different ext-BCM-based regressors. Panels (a) and (e) display the ground-truth simulations obtained directly from the ext-BCM ODE solver. RF: Random forest. NN: Neural Network. PR: Polynomial regressor.

On the other hand, regarding the capacity of the three regressors as a model of the plant in a trajectory tracking scheme, Figure 4 presents the tracking results for pitch glide (a) and loudness glide (b). The regressors effectively follow the reference trajectory, depicted as a black dashed line.

**FIG. 4.**
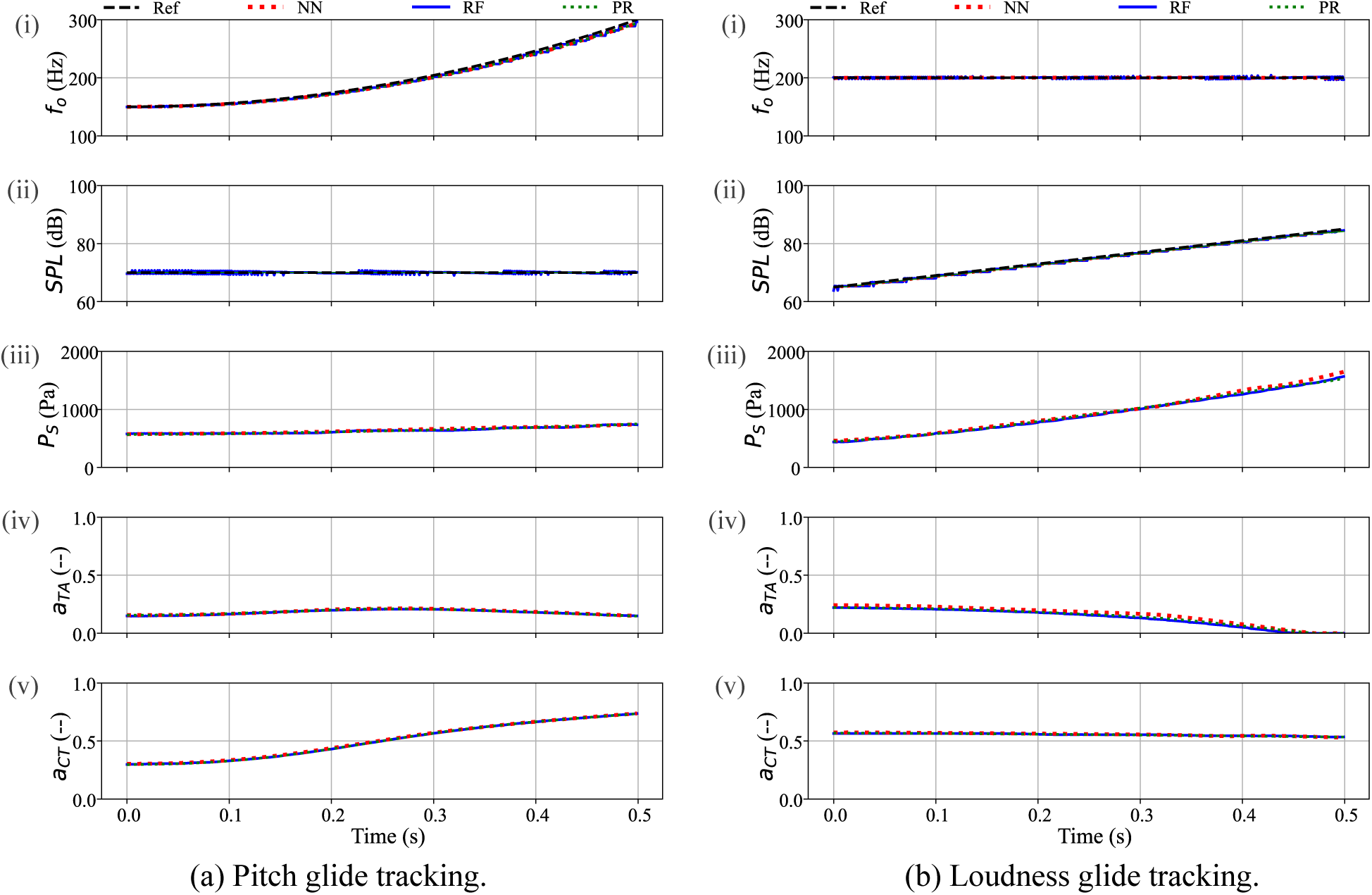
Tracking glides for ext-BCM-based regressors. Column (a) corresponds to a pitch-glide gesture, while column (b) corresponds to a loudness-glide gesture. Target/Output signals: (i) f*_o_* and (ii) SPL. Input signals: (iii) subglottal pressure P*_S_*, (iv) TA activation a*_T A_*, and (v) CT activation a*_CT_*. The black dashed line represents the reference gesture. These reference glides are mathematically synthesized kinematic targets used to evaluate tracking stability, not recordings from human subjects. The red dashed line shows the performance of a Neural-Networks regressor, the blue solid line correspond to a Random Forest regressor, and the green dotted line corresponds a Polynomial regressor.

Regarding f*_o_* and SPL references (i-ii), all regressors exhibit a smooth tracking behavior. However, a step-like pattern can be observed in the RF regressor, especially in the SPL trajectories, because estimates values are based on discrete samples from the training dataset. In contrast, the NN and PR models generate continuous outputs. Despite these differences, all three regressors result in the same motor control input behavior (iii-v).

For pitch glide, an increase in CT activation and a slight decrease in TA activation are observed, aligning with their roles in the regulation of VFs tension (Chhetri and Park, 2016; Yin and Zhang, 2013). P*_S_* tends to remain constant, as a fixed SPL was imposed.

In contrast, in the loudness glide, the a*_CT_* tends to remain constant while a*_T A_* decreases, which diverges from previous electromyography evidence, which typically reports an increase in a*_T A_* together with elevated P*_S_* during intensity rises (Baker *et al*., 2001). This discrepancy may reflect a limitation of the ext-BCM model in capturing the adductory adjustments that normally accompany loudness control. The observed SPL rise, driven by P*_S_*, nevertheless aligns with its well-known strong correlation with subglottal pressure (Chhetri and Park, 2016) and with the fact that SPL is a logarithmic measure of sound pressure.

### B. Evaluation of machine learning regressors for forward mapping and reference tracking for the TBCM dataset

Before evaluating the regressors on the TBCM dataset, we performed a preliminary assessment of the impact of dataset size (N) on predictive accuracy. This analysis, illustrated in Figure 5, serves to identify the minimum data requirements for learning the complex, non-linear mapping of laryngeal biomechanics from scratch. As observed in the log-log plots for both MAE and RMSE, performance remains stable and high for large datasets (N > 10, 000). However, a significant exponential degradation in accuracy occurs when the training size drops below N = 1, 000, particularly for f*_o_* predictions. This suggests that without sufficient empirical evidence, standard ML architectures struggle to capture the underlying physics and bifurcations of the biomechanical system. These results establish a critical baseline: while N ≈ 50, 000 (TBCM size) is sufficient for high-fidelity modeling, clinical scenarios with N < 100 samples would strictly require TL strategies rather than training from scratch.

**FIG. 5.**
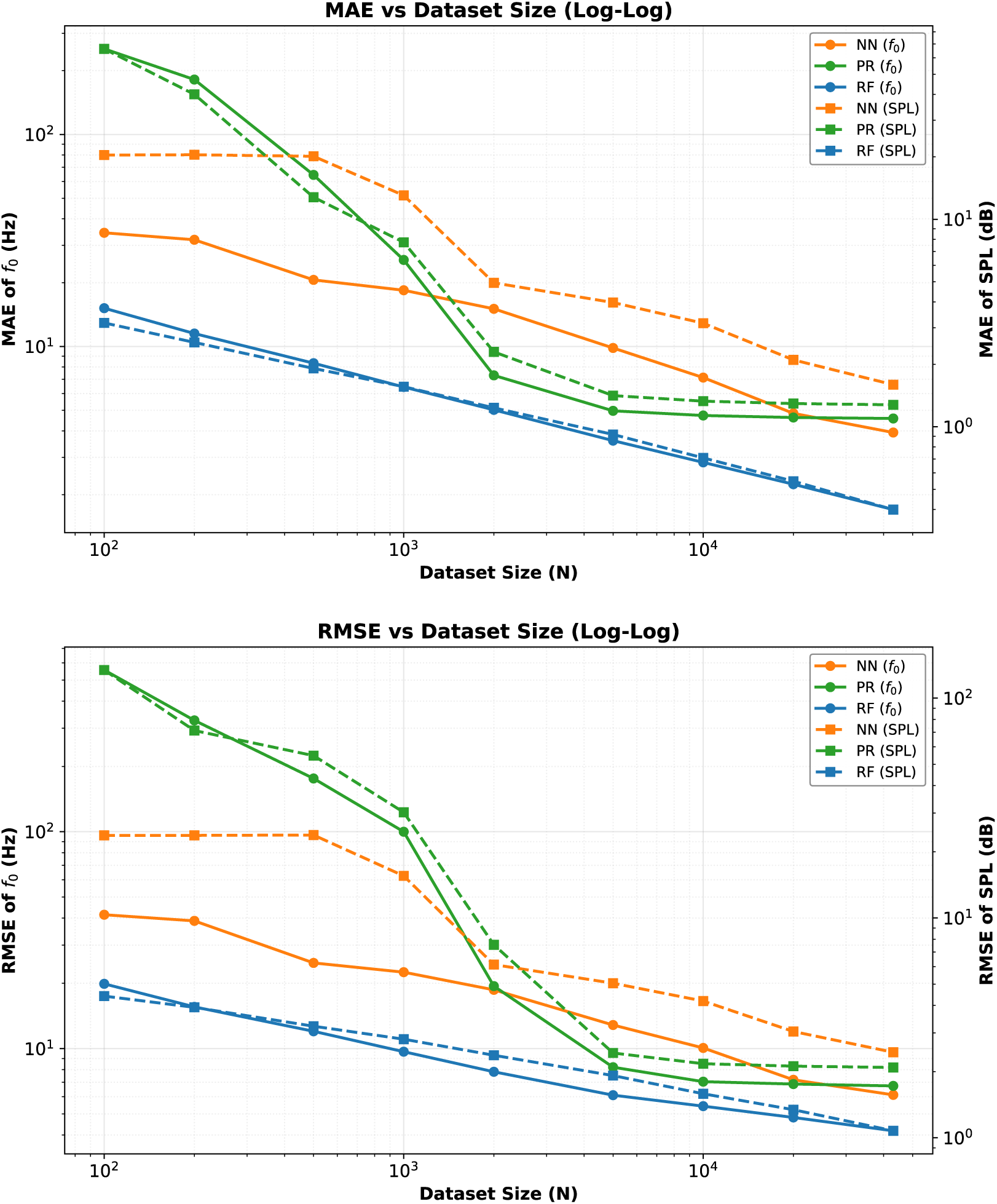
Effect of training dataset size (N) on the predictive accuracy of the ML regressors. The top panel displays Mean Absolute Error (MAE), and the bottom panel displays Root Mean Square Error (RMSE) for f*_o_* (solid lines, left axis) and SPL (dashed lines, right axis) on a log-log scale. A significant degradation in performance is observed for N < 1, 000.

Building upon the dataset size analysis (Figure 5), this section evaluates the regressors on the TBCM dataset. With approximately 50,000 samples, this dataset size remains well within the high-accuracy regime identified in our preliminary assessment, ensuring that the results primarily reflect the regressors’ capability to adapt to a more complex laryngeal representation (i.e., triangular geometry and detailed muscle constitutive laws (Titze and Hunter, 2007)). This setup provides a robust foundational scenario to test whether the optimized architectures can maintain their predictive performance when the underlying biomechanical mapping changes.

This analysis evaluates the combined effect of training with a more complex laryngeal representation (TBCM) and a substantially smaller dataset. The advantage of this design is that it provides a first synthetic scenario resembling real-world applications, where models must deal with both increased physiological detail and limited data availability. The drawback, however, is that it becomes difficult to fully disentangle the contributions of model complexity and dataset size to performance changes.

To assess this adaptability, we re-trained the regressors on the TBCM dataset while keeping the optimized structures identified in the previous section. Table IV presents the results, showing R^2^ values above 0.94 in all cases. While there is a slight increase in error compared to the ext-BCM results, the performance remains high, confirming that the models successfully captured the added complexity of the TBCM representation. The observed trend remains: RF attains the lowest errors (1.7 Hz for f*_o_* and 0.4 dB for SPL), while the performance gap with PR and NN has narrowed, with the latter showing errors of approximately 4–6 Hz for f*_o_* and 1.3 dB for SPL.

**TABLE IV.** Results for optimal regressors applied to TBCM dataset, using Random Forest (RF), Neural Network (NN) and Polynomial Regressor (PR).

| | $f_o$ | | | $SPL$ | | |
| --- | --- | --- | --- | --- | --- | --- |
| | $R^2$ | MAE (Hz) | RMSE (Hz) | $R^2$ | MAE (dB) | RMSE (dB) |
| RF | 0.996 | 1.733 | 4.279 | 0.988 | 0.401 | 1.038 |
| NN | 0.993 | 3.668 | 5.676 | 0.948 | 1.346 | 2.203 |
| PR | 0.991 | 4.566 | 6.678 | 0.955 | 1.274 | 2.049 |

Regarding the computational time for training, since the regressor structures remain unchanged, the execution times are the same as reported in Figure 2. However, a notable change was observed in the size of the RF model: Due to the smaller training dataset, its size was reduced to 140 MB, while the sizes of the other two regressors remained unchanged.

Heatmaps for f*_o_* and SPL (Figures 6) illustrate the differences in phonatory representation between the regressors, with respect to the simulated data. The TBCM model introduces more complex and non-linear biomechanical interactions. While such richness provides an advantage for ML regressors to capture detailed relationships, it also increases the challenge, as reflected in the observed performance differences compared to ext-BCM. A distinct pattern was observed in the relationship between f*_o_* and muscle activation, in comparison with the ext-BCM heatmaps. The training data size also affected the heatmap shapes between the regressors. SPL proved more challenging to reproduce, particularly for RF (Figure 6 f), where squared patterns emerged. These may reflect the hierarchical partitioning intrinsic to tree-based models, but could also be partially influenced by the discretization of the training grid. In contrast, PR (Figure 6 h) produced smoother patterns due to its polynomial nature, while NN (Figure 6 g) generated SPL representations more consistent with the original simulated data.

**FIG. 6.**
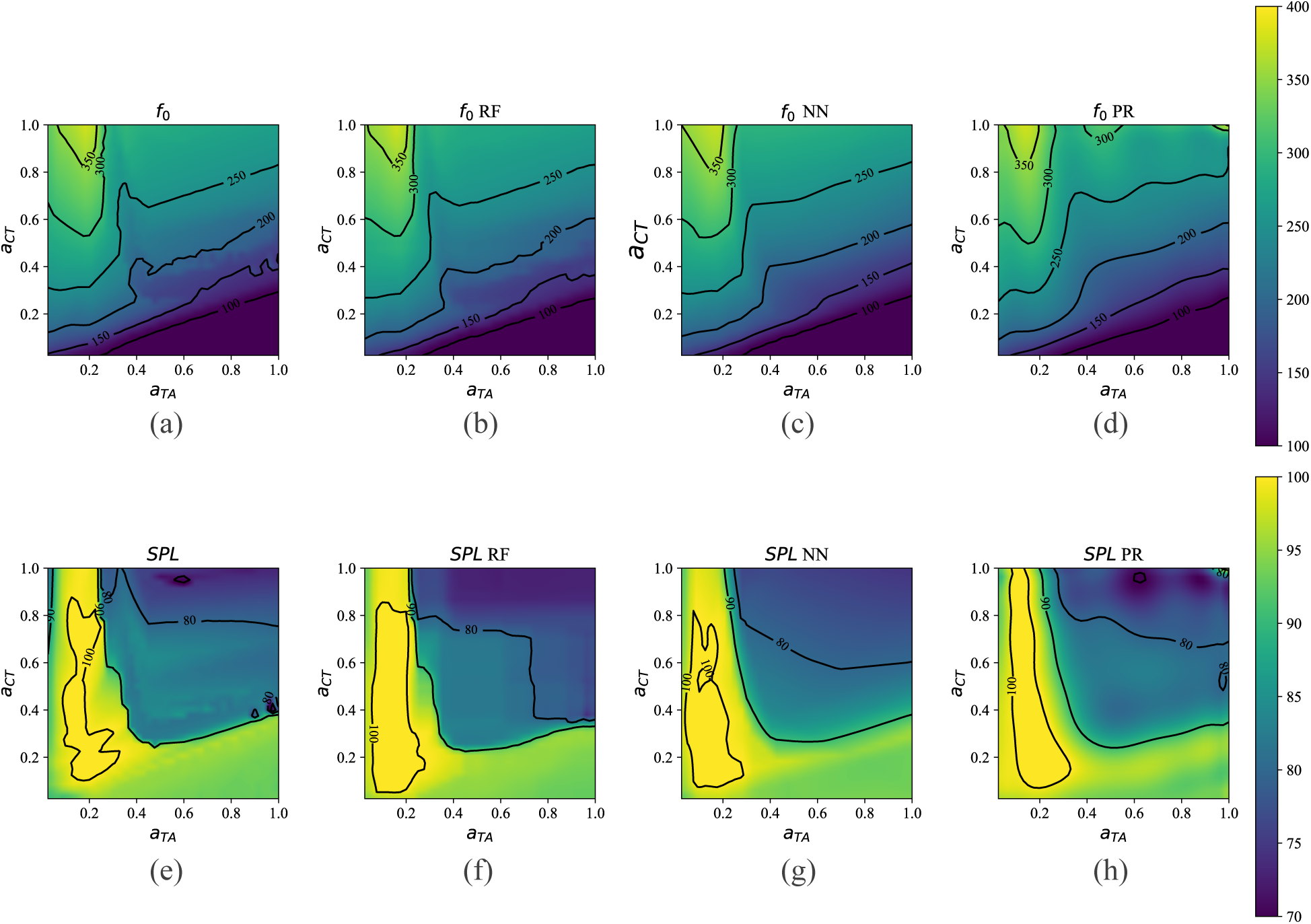
MAP of f*_o_* (a-d) and SPL (e-h) for the different TBCM-based regressors. Panels (a) and (e) display the ground-truth simulations obtained directly from the TBCM ODE solver. RF: Random forest. NN: Neural Network. PR: Polynomial regressor.

The trajectory tracking results for the TBCM-trained regressors are presented in Figure 7. While the regressors are capable of following the gesture trajectories, the RF model exhibits some instability.

**FIG. 7.**
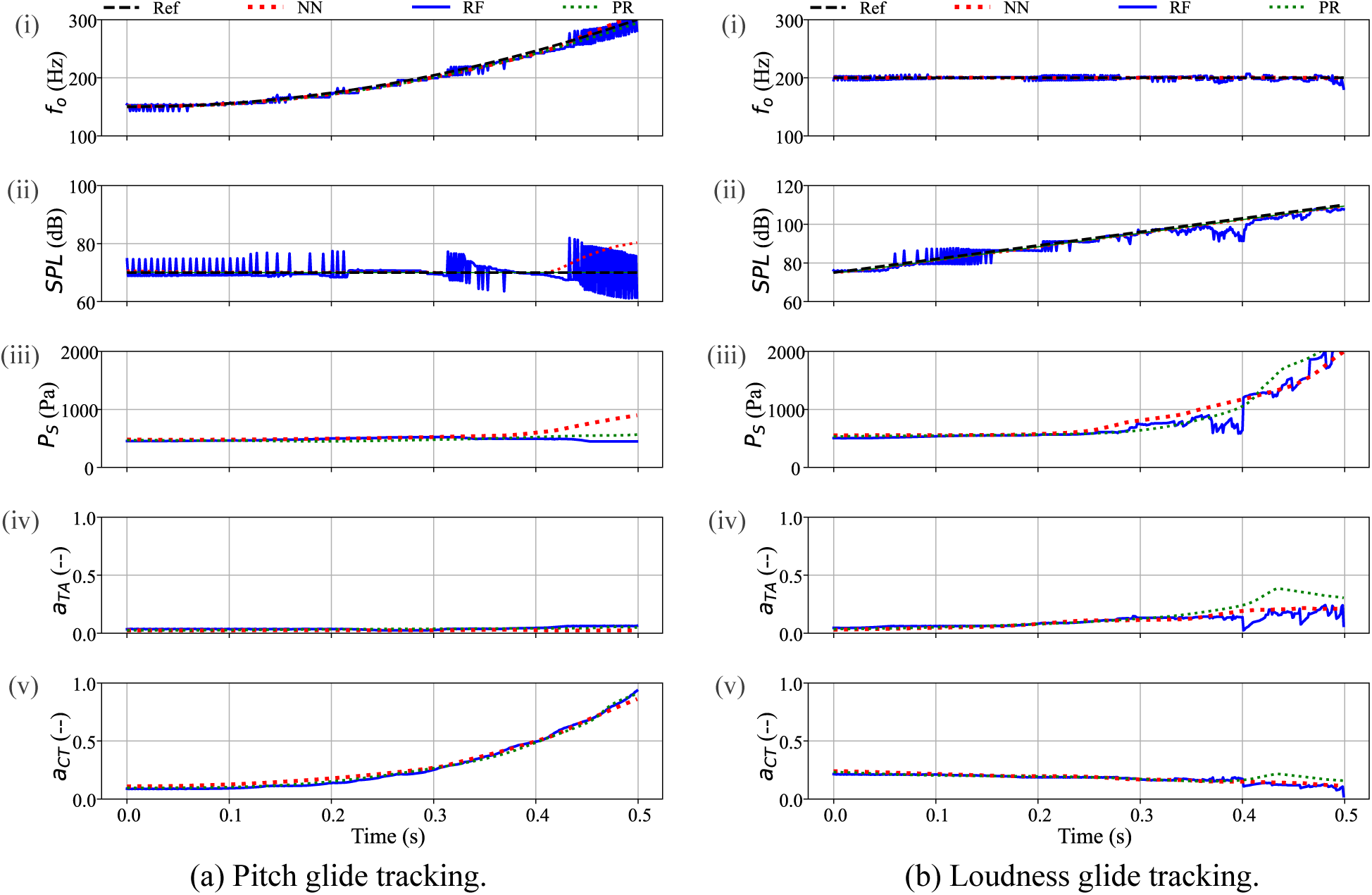
Tracking glides for TBCM-based regressors. Column (a) corresponds to a pitch-glide gesture, while column (b) corresponds to a loudness-glide gesture. Target/Output signals: (i) f*_o_* and (ii) SPL. Input signals: (iii) subglottal pressure P*_S_*, (iv) TA activation a*_T A_*, and (v) CT activation a*_CT_*. The black dashed line represents the reference gesture. These reference glides are mathematically synthesized kinematic targets used to evaluate tracking stability. The red dashed line shows the performance of a Neural-Networks regressor, the blue solid line correspond to a Random Forest regressor, and the green dotted line corresponds a Polynomial regressor.

For the pitch glide (a), RF struggles to maintain stable SPL (ii), showing oscillatory behavior at certain time instants, with fewer difficulties in tracking the increase in f*_o_* (i). Regarding the control signal (iii-v), a similar pattern to that obtained for ext-BCM is observed, where the f*_o_* increase is primarily driven by increasing CT activation. However, discrepancies between the regressors are noticeable at the end of the gesture, where the target f*_o_* is at its upper limit around 300 Hz. RF exhibits greater fluctuations, while NN increases P*_S_*, which is consistent with an increase in SPL. For the loudness glide (b), the increase in SPL is primarily driven by a gradual increase in P*_S_*. The P*_S_* trajectory (iii) increases between the three regressors as time evolves, and muscle activations tend to remain constant, although differences between the regressors emerge toward the end of the gesture, with few increases in TA and decreases in CT. This behavior is expected, as 110 dB is at the upper extreme of the range of the TBCM model.

## IV. DISCUSSION

The comparative evaluation of ML architectures demonstrated that tree-based models, specifically Random Forest (RF), achieved superior performance on large synthetic datasets such as the ext-BCM (N ≈ 800, 000). RF reached R^2^ values of 0.999, with mean absolute errors significantly lower than those of Neural Networks (NN) and Polynomial Regression (PR). Specifically, NN and PR exhibited errors for f*_o_* that were more than three times higher than those of RF, and for SPL, the errors were between 8 and 15 times greater. These results confirm that, given sufficient empirical evidence, ensemble tree methods excel at capturing the high-dimensional, tabular nonlinearities inherent in laryngeal biomechanics.

However, the performance landscape shifts as the dataset size decreases. Our results with the TBCM dataset (N ≈ 50, 000) show that the performance gap between regressors tends to narrow. This behavior is further elucidated by the dataset size analysis (Figure 5), which identifies a critical threshold at approximately N = 1, 000 samples. Below this point, a significant exponential degradation in accuracy and tracking stability occurs for all architectures trained from scratch. This threshold represents a fundamental limitation: without sufficient data to map the complex nonlinear bifurcations and transitions of the vocal fold system, standard ML architectures cannot reliably learn the underlying physics. Consequently, while N ≈ 50, 000 is sufficient to capture detailed biomechanical interaction, the typically limited nature of clinical datasets (N < 100) makes training complex forward models from scratch an impractical approach for subject-specific applications.

The choice of an appropriate regressor for laryngeal control therefore involves a critical trade-off between predictive accuracy, model complexity, and computational cost (Choo and Chang, 2022). While RF provides the highest accuracy, it does so at the cost of significantly higher memory demands due to the storage of numerous decision nodes. In contrast, NN and PR require only the storage of weight matrices, resulting in a much smaller memory footprint. Crucially, in iterative trajectory tracking tasks where the plant must be evaluated repeatedly to compute numerical derivatives, execution time becomes as vital as accuracy. In this regard, PR and NN offer a clear advantage; for instance, the PR’s execution time (≈ 2 ms) is orders of magnitude faster than the direct numerical integration of the underlying ODEs (≈ 6 s), making real-time control viable.

This data-size limitation, combined with the need for computational efficiency, strongly motivates the transition toward TL strategies. While personalizing traditional ODE-based models requires solving highly complex, ill-posed inverse optimization problems to adjust hidden anatomical hyperparameters (e.g., resting VFs length, tissue stiffness, or muscle constants), an ML surrogate provides a flexible architecture perfectly suited for data-driven fine-tuning. The foundational ML regressors established in this study can act as “base models” that have already learned the fundamental physical rules of phonation from large synthetic datasets. These models can then be efficiently adapted to individual subjects using small clinical datasets without the need for training from scratch.

For NN architectures, this fine-tuning approach is straightforward, involving the adjustment of specific layers to capture subject-specific nuances (Ibarra *et al*., 2025). For RF, similar strategies can involve building ensembles across datasets (Gu *et al*., 2022) or refining branch weights based on patient data (Segev *et al*., 2016). PR can also benefit from two-stage task-transferring methods, such as Lasso-type regularization or gradient-descent fine-tuning (Li *et al*., 2022; Obst *et al*., 2022). Furthermore, incorporating probabilistic regressors, such as Gaussian Processes or Kriging-based models, represents a highly valuable direction for future TL applications. These models provide a principled framework for uncertainty quantification and have demonstrated strong performance in data-scarce scenarios by leveraging correlations between data-rich (synthetic) and data-limited (clinical) domains through multi-task or mixture-of-experts formulations (Eleftheriadis *et al*., 2017; Iapteff *et al*., 2021; Yao *et al*., 2025).

In the tracking experiments, the observed trends with the TBCM-based regressors successfully align with established laryngeal physiology. The models accurately reproduced the monotonic increase of f*_o_* with CT activation and the high sensitivity of SPL to subglottal pressure (Martínez *et al*., 2025). Importantly, the surrogates also captured complex antagonistic relationships and the non-monotonic effect of TA activation on f*_o_*, which acts as either a pitch raiser or lowerer depending on the specific region of the CT-TA activation space. While slight mismatches in adjustment magnitudes were observed, these likely reflect simplifications in the underlying lumped-element models or the existence of multiple inverse solutions in the Jacobian-based approach, which tends to favor minimum-norm solutions (Chhetri and Park, 2016; Chung *et al*., 2024).

Several limitations of the present work should be noted. The evaluation used 50-ms analysis windows, providing a quasi-stationary approximation that may not fully capture rapid phonatory transients (Weerathunge *et al*., 2022). Furthermore, the regressors were trained within predefined parameter ranges, limiting their reliability in extrapolation (e.g., laryngeal configurations with a*_LC_* > 0.6 or other conditions not represented in the training set). The models also inherit the simplifications of the underlying lumped-element solvers, such as idealized geometries and linear material properties. Additionally, the training relies on simulated data generated under idealized conditions. Consequently, the current datasets lack the acoustic variance or pathological perturbations (e.g., tremor, jitter) typically found in clinical signals. Future work must incorporate clinical datasets to validate robustness against these real-world variances. Moreover, only two lumped-element VFs models were considered; additional configurations that represent alternative anatomical or pathological conditions (Parra *et al*., 2024; Vahabzadeh-Hagh *et al*., 2018) could provide a broader validation space and strengthen the applicability of the framework. Finally, the challenge of non-unique inverse solutions in tracking can be addressed by incorporating null-space formulations to introduce additional physiological constraints (Chen and Walker, 1993). Another promising path is the direct integration of ML surrogates into the inverse control system itself (Palaparthi *et al*., 2024), building upon previous efforts in laryngeal parameter estimation (Donhauser *et al*., 2024; Ibarra *et al*., 2025, 2021; Zhang, 2020).

## V. CONCLUSION

The results of this study demonstrate that feature-driven ML regressors are highly effective as surrogate forward models for laryngeal biomechanical control simulations. By replacing computationally restrictive numerical ODE solvers, these ML plants reduce execution times from seconds to milliseconds, enabling real-time evaluations within dynamic closed-loop simulations. Among the evaluated methods, tree-based approaches, particularly RF, achieved the highest predictive accuracy for f*_o_* and SPL. However, this advantage came at the cost of higher memory demands and less stable trajectories during iterative tracking tasks, where PR and NN provided smoother, continuous control signals and faster computation.

Model performance was found to be highly sensitive to both the complexity of the underlying biomechanical representation and the size of the training dataset. Crucially, the evaluation of data scarcity revealed a critical degradation threshold around N = 1, 000 samples. Below this limit, the lack of sufficient empirical evidence prevents standard ML architectures from reliably learning the complex nonlinear bifurcations of the VFs system from scratch.

Consequently, the optimal choice of ML regressor involves a strict trade-off between predictive performance, memory capacity, and execution speed. When large, complex datasets are available and memory is not a constraint, RF provides the best static accuracy. Conversely, in scenarios with limited data or where smooth dynamic tracking is prioritized, PR and NN become more competitive alternatives due to their compact analytical structures and interpolation capabilities.

These findings validate the feasibility of replacing rigid differential equation solvers with fast, adaptable ML forward models. By establishing this robust baseline and clearly identifying the data-size limitations of training from scratch, the present work provides the necessary physical and computational justification for future TL strategies. Ultimately, this lays the mandatory groundwork for integrating subject-specific laryngeal biomechanics into clinical and speech motor control frameworks without relying on complex, ill-posed inverse ODE optimization.

## ACKNOWLEDGMENTS

This research was supported by the National Institutes of Health (NIH) National Institute on Deafness and Other Communication Disorders grant P50 DC015446, ANID grants Becas de Doctorado Nacional 21202490 and 21240471, FONDECYT 1230828, 3250844 and 3260566, and BASAL AFB240002. The content is solely the responsibility of the authors and does not necessarily represent the official views of the NIH.

## DATA AVAILABILITY

Upon acceptance, the simulation datasets, trained surrogate models, and training/evaluation scripts will be made publicly available in a GitHub repository (github.com/jealpape/ML-Regressors-for-Laryngeal-Biomechanical-Control). The repository will include instructions and a containerized environment to reproduce the experiments.

## REFERENCES

Alzamendi, G. A., Peterson, S. D., Erath, B. D., Hillman, R. E., and Zañartu, M. (2022). “Triangular body-cover model of the vocal folds with coordinated activation of the five intrinsic laryngeal muscles,” The Journal of the Acoustical Society of America 151(1), 17–30.

Baker, K. K., Ramig, L. O., Sapir, S., Luschei, E. S., and Smith, M. E. (2001). “Control of vocal loudness in young and old adults,”.

Chen, Y.-C., and Walker, I. D. (1993). “A consistent null-space based approach to inverse kinematics of redundant robots,” in [1993] Proceedings IEEE International Conference on Robotics and Automation, IEEE, pp. 374–381.

Chhetri, D. K., and Neubauer, J. (2015). “Differential roles for the thyroarytenoid and lateral cricoarytenoid muscles in phonation,” The Laryngoscope 125(12), 2772–2777, doi: 10.1002/lary.25480.

Chhetri, D. K., and Park, S. J. (2016). “Interactions of subglottal pressure and neuromuscular activation on fundamental frequency and intensity,” The Laryngoscope 126(5), 1123–1130, https://onlinelibrary.wiley.com/doi/abs/10.1002/lary.25550, doi: 10.1002/lary.25550.

Choo, Y. J., and Chang, M. C. (2022). “Use of machine learning in stroke rehabilitation: a narrative review,” Brain & Neurorehabilitation 15(3), e26.

Chung, H. R., Lee, Y., Reddy, N. K., Zhang, Z., and Chhetri, D. K. (2024). “Effects of thyroarytenoid activation induced vibratory asymmetry on voice acoustics and perception,” The Laryngoscope 134(3), 1327–1332.

Donhauser, J., Tur, B., and Döllinger, M. (2024). “Neural network-based estimation of biomechanical vocal fold parameters,” Frontiers in Physiology 15, 1282574.

Eleftheriadis, S., Rudovic, O., Deisenroth, M. P., and Pantic, M. (2017). “Gaussian process domain experts for modeling of facial affect,” IEEE transactions on image processing 26(10), 4697–4711.

Galindo, G. E., Peterson, S. D., Erath, B. D., Castro, C., Hillman, R. E., and Zañartu, M. (2017). “Modeling the pathophysiology of phonotraumatic vocal hyperfunction with a triangular glottal model of the vocal folds,” Journal of Speech, Language, and Hearing Research 60(9), 2452–2471.

Gomez, P., Fernández-Baillo, R., Nieto, A., Díaz, F., Fernández-Camacho, F., Rodellar, V., Álvarez Marquina, A., and Martínez-Olalla, R. (2007). “Evaluation of voice pathology based on the estimation of vocal fold biomechanical parameters,” Journal of voice: official journal of the Voice Foundation 21, 450–76, doi: 10.1016/j.jvoice.2006.01.008.

Grinsztajn, L., Oyallon, E., and Varoquaux, G. (2022). “Why do tree-based models still outperform deep learning on typical tabular data?,” in NeurIPS Datasets and Benchmarks Track.

Gu, T., Han, Y., and Duan, R. (2022). “A transfer learning approach based on random forest with application to breast cancer prediction in underrepresented populations,” in Biocomputing 2023, WORLD SCIENTIFIC.

Guenther, F. H. (2016). Neural control of speech (Mit Press).

Hillman, R. E., Stepp, C. E., Stan, J. H. V., Zañartu, M., and Mehta, D. D. (2020). “An updated theoretical framework for vocal hyperfunction,” American Journal of Speech-Language Pathology 29(4), 2254–2260, https://pubs.asha.org/doi/abs/10.1044/2020_AJSLP-20-00104, doi: 10.1044/2020_AJSLP-20-00104.

Hunt, K. J., Sbarbaro, D., Żbikowski, R., and Gawthrop, P. J. (1992). “Neural networks for control systems—a survey,” Automatica 28(6), 1083–1112.

Iapteff, L., Jacques, J., Rolland, M., and Celse, B. (2021). “Reducing the number of experiments required for modelling the hydrocracking process with kriging through bayesian transfer learning,” Journal of the Royal Statistical Society Series C: Applied Statistics 70(5), 1344–1364.

Ibarra, E. J., Arias-Londoño, J. D., Godino-Llorente, J. I., Mehta, D. D., and Zañartu, M. (2025). “Subject-specific modeling by domain adaptation for the estimation of subglottal pressure from neck-surface acceleration signals,” Biomedical Signal Processing and Control 106, 107681.

Ibarra, E. J., Parra, J. A., Alzamendi, G. A., Cortés, J. P., Espinoza, V. M., Mehta, D. D., Hillman, R. E., and Zañartu, M. (2021). “Estimation of subglottal pressure, vocal fold collision pressure, and intrinsic laryngeal muscle activation from neck-surface vibration using a neural network framework and a voice production model,” Frontiers in Physiology 12.

Kinahan, S. P., Liss, J. M., and Berisha, V. (2023). “Torchdiva: An extensible computational model of speech production built on an open-source machine learning library,” PLOS ONE 18(2), e0281306.

Kröger, B. J., Kannampuzha, J., and Neuschaefer-Rube, C. (2009). “Towards a neurocomputational model of speech production and perception,” Speech Communication 51(9), 793–809.

Li, S., Cai, T. T., and Li, H. (2022). “Transfer learning for high-dimensional linear regression: Prediction, estimation and minimax optimality,” Journal of the Royal Statistical Society Series B: Statistical Methodology 84(1), 149–173.

Marks, K. L., Lin, J. Z., Burns, J. A., Hron, T. A., Hillman, R. E., and Mehta, D. D. (2020). “Estimation of subglottal pressure from neck surface vibration in patients with voice disorders,” Journal of Speech, Language, and Hearing Research 63(7), 2202–2218, doi: 10.1044/2020_JSLHR-19-00409.

Martínez, J. D., Ibarra, E. J., Parra, J. A., Mehta, D. D., Heaton, J. T., Hillman, R. E., Plocienniczak, M. J., Cooper, J. C., and Zañartu, M. (2025). “Toward acoustic-based normalization of laryngeal emg for improved interspeaker consistency in muscle-to-acoustic mapping,” Journal of Voice.

Obst, D., Ghattas, B., Claudel, S., Cugliari, J., Goude, Y., and Oppenheim, G. (2022). “Improved linear regression prediction by transfer learning,” Computational Statistics & Data Analysis 174, 107499.

Palaparthi, A., Alluri, R. K., and Titze, I. R. (2024). “Deep learning for neuromuscular control of vocal source for voice production,” Applied Sciences 14(2), 769.

Parra, J. A., Calvache, C., Alzamendi, G. A., Ibarra, E. J., Soláque, L., Peterson, S. D., and Zañartu, M. (2024). “Asymmetric triangular body-cover model of the vocal folds with bilateral intrinsic muscle activation,” The Journal of the Acoustical Society of America 156(2), 939–953.

Parrell, B., Lammert, A. C., Ciccarelli, G., and Quatieri, T. F. (2019). “Current models of speech motor control: A control-theoretic overview of architectures and properties,” The Journal of the Acoustical Society of America 145(3), 1456–1481.

Pedregosa, F., Varoquaux, G., Gramfort, A., Michel, V., Thirion, B., Grisel, O., Blondel, M., Prettenhofer, P., Weiss, R., Dubourg, V., Vanderplas, J., Passos, A., Cournapeau, D., Brucher, M., Perrot, M., and Duchesnay, E. (2011). “Scikit-learn: Machine learning in Python,” Journal of Machine Learning Research 12, 2825–2830.

Perrier, P., Ma, L., and Payan, Y. (2006). “Modeling the production of vcv sequences via the inversion of a biomechanical model of the tongue,” arXiv preprint physics/0610170.

Rudin, C. (2019). “Stop explaining black box machine learning models for high stakes decisions and use interpretable models instead,” Nature Machine Intelligence 1(5), 206–215.

Saltzman, E. L., and Munhall, K. G. (1989). “A dynamical approach to gestural patterning in speech production,” Ecological psychology 1(4), 333–382.

Segev, N., Harel, M., Mannor, S., Crammer, K., and El-Yaniv, R. (2016). “Learn on source, refine on target: A model transfer learning framework with random forests,” IEEE transactions on pattern analysis and machine intelligence 39(9), 1811–1824.

Story, B. H. (2008). “Comparison of magnetic resonance imaging-based vocal tract area functions obtained from the same speaker in 1994 and 2002,” The Journal of the Acoustical Society of America 123(1), 327–335.

Story, B. H., and Titze, I. R. (1995). “Voice simulation with a body-cover model of the vocal folds,” Journal of the Acoustical Society of America 97(2), 1249–1260.

Titze, I. (2000). Principles of Voice Production (National Center for Voice and Speech), https://books.google.cl/books?id=ytAeAQAAMAAJ.

Titze, I. R., and Hunter, E. J. (2007). “A two-dimensional biomechanical model of vocal fold posturing,” The Journal of the Acoustical Society of America 121(4), 2254.

Titze, I. R., and Story, B. H. (2002). “Rules for controlling low-dimensional vocal fold models with muscle activation,” The Journal of the Acoustical Society of America 112(3), 1064–1076.

Tourville, J. A., and Guenther, F. H. (2011). “The diva model: A neural theory of speech acquisition and production,” Language and cognitive processes 26(7), 952–981.

Vahabzadeh-Hagh, A. M., Zhang, Z., and Chhetri, D. K. (2018). “Hirano’s cover–body model and its unique laryngeal postures revisited,” The Laryngoscope 128(6), 1412–1418.

Weerathunge, H. R., Alzamendi, G. A., Cler, G. J., Guenther, F. H., Stepp, C. E., and Zañartu, M. (2022). “Ladiva: A neurocomputational model providing laryngeal motor control for speech acquisition and production,” PLOS Computational Biology 18(6), e1010159.

Yao, J., Wu, J., Li, Y., and Wang, C. (2025). “Transfer learning of stochastic kriging for individualized prediction,” IEEE transactions on pattern analysis and machine intelligence.

Yin, J., and Zhang, Z. (2013). “The influence of thyroarytenoid and cricothyroid muscle activation on vocal fold stiffness and eigenfrequencies,” The Journal of the Acoustical Society of America 133(5), 2972–2983.

Zañartu, M., Galindo, G. E., Erath, B. D., Peterson, S. D., Wodicka, G. R., and Hillman, R. E. (2014). “Modeling the effects of a posterior glottal opening on vocal fold dynamics with implications for vocal hyperfunction,” The Journal of the Acoustical Society of America 136(6), 3262–3271.

Zhang, Z. (2016). “Mechanics of human voice production and control,” The Journal of the Acoustical Society of America 140(4), 2614–2635, doi: 10.1121/1.4964509.

Zhang, Z. (2020). “Estimation of vocal fold physiology from voice acoustics using machine learning,” The Journal of the Acoustical Society of America 147(3), EL264–EL270.

